# Genotype by environment interactions for the shade avoidance syndrome in the plant *Antirrhinum majus*, the snapdragon

**DOI:** 10.1101/331033

**Authors:** Mathilde Mousset, Sara Marin, Juliette Archambeau, Christel Blot, Vincent Bonhomme, Laura Garaud, Benoit Pujol

## Abstract

A classical example of phenotypic plasticity in plants is the set of trait changes in response to shade, *i.e.* the shade avoidance syndrome. There is widespread evidence that plants in low light conditions often avoid shade by growing taller or by increasing their photosynthetic efficiency. This plastic response is expected to have evolved in response to selection in several species, yet there is limited evidence for its genetic variation within populations, which is required for any evolutionary response to selection. In this study, we investigated the shade avoidance syndrome in snapdragon plants (*Antirrhinum majus*) by using a common garden approach on four natural populations from the Mediterranean region. Our results showed that, in the four populations, individual plants reacted strongly to the presence of shade by growing longer shoots, longer internodes, and increasing their specific leaf area. Our results also revealed genetic variation for the plastic response within these populations, as well as few genetic constraints to its evolution. Our findings imply that the plastic response to shade has the potential to evolve in response to selection in natural populations of *A. majus*.

## Introduction

Environmental heterogeneity is the norm rather than the exception in nature. As a consequence, organisms often experience variation in their environment across space and time. In this case, theory predicts that phenotypic plasticity (the capacity of one genotype to express different trait values in different environments) is likely to evolve (Via and Lande, 1985; Scheiner, 1993, 2013; Gavrilets and Scheiner, 1993). The capacity of a population to evolve different levels of phenotypic plasticity is conditioned, however, by the presence of genetic variation for the plastic response. There is a large body of evidence showing a genetic basis to phenotypic plasticity for many traits (e.g. Beldade *et al.*, 2011). A variety of approaches can be used to test for this genetic component (Pigliucci, 2005). One classical method involves testing populations for gene-by-environment (GxE) interactions, by investigating how identical or related genotypes express different phenotypes in responses to a change in their environmental conditions. A genetic component to plasticity is therefore evident if related individuals express a more similar plastic response than non-related individuals. Here, we conducted a common garden experiment and used quantitative genetic models to evaluate the evolutionary potential of the plastic response to shade of four natural populations of snapdragon plants (*Antirrhinum majus* L.).

A classic example of phenotypic plasticity is the response of plants when light intensity and spectrum are heterogeneous (e.g. Smith, 1982; Schmitt and Wulff, 1993; Schmitt *et al.*, 1995, 1999). In many species, it has been documented that shade triggers a suite of developmental, physiological, and phenological trait modifications. These modifications can include the increased growth of the shoot (often through internode elongation), petiole or hypocotyl (Smith, 1982; Ballaré *et al.*, 1991; Ballaré, 1999; Pierik *et al.*, 2004; Pujol *et al.*, 2005; Bell and Galloway, 2007; Franklin, 2008), modification of the position of leaves (Pierik *et al.*, 2003), increased apical dominance (Smith, 1982; Smith and Whitelam, 1997), reduction of branching or tillering (Casal et al., 1986, Smith, 1982; Schmitt and Wulff 1993), earlier flowering (Smith, 1982; Schmitt and Wulff, 1993), and a greater leaf area to biomass ratio (increase of specific leaf area, (Lewandowska and Jarvis, 1977; Dong, 1995; Evans and Poorter, 2001). Collectively, these trait modifications characterise the shade avoidance syndrome. The molecular mechanisms underlying this syndrome are increasingly well described in mutants or inbred lines (e.g. Ballaré *et al.*, 1987, 1990; Schmitt *et al.*, 1995; Ballaré and Pierik, 2017), and involve the action of photoreceptors and phytohormones.

The shade avoidance syndrome in plants is ecologically significant in many species. It is acknowledged as a functional mechanism that increases the ability of plants to reach or exploit light in the presence of competition. Stem elongation, for example, can result in longer shoots with the top leaves reaching above the canopy of competitors. The presence of this functional response in natural populations implies that past evolution has shaped the plant plasticity to light conditions. However, to have contemporary evolution requires both selection pressures and heritable variation. While the shade avoidance syndrome is often considered to be adaptive (Schmitt *et al.*, 2003), few studies demonstrated its association with a fitness advantage (van Kleunen and Fischer, 2001; Schmitt *et al.*, 2003; Bell and Galloway, 2007). Furthermore, much less is known about the genetic variation of the shade avoidance syndrome in natural populations, especially in regard to the extent of growing knowledge about its molecular mechanisms. Two notable exceptions are the work of Bell & Galloway (2007) on *Geranium,* and a series of studies from Donohue and colleagues in field-derived inbred lines of *Impatiensis capensis* (e.g. Donohue and Schmitt, 1999; Donohue *et al.*, 2000; Huber *et al.*, 2004). Most data on the heritable genetic variation for the magnitude of the plastic response of plants to shade is based, however, on inbred lines and variation between specific mutants (e.g. Schmitt *et al.*, 1995; Donohue *et al.*, 2000; Botto and Smith, 2002; Huber *et al.*, 2004).

In this article, we investigated the presence of the shade avoidance syndrome in individuals of four natural populations of *A. majus* in a common garden experiment. *A. majus* grows in Mediterranean garrigues (dry scrubland comprised of a mixture of shrub and herb species) and Pyrenean mountain habitats, where there is variation in the density of plants and competition for light at both spatial and temporal scales. We therefore expected natural *A. majus* populations to exhibit a suite of plastic responses that are typical of the shade avoidance syndrome. We used a quantitative genetic approach to test for 1) the presence of a shade avoidance syndrome in these four natural populations and 2) genetic variation for the shade avoidance response in these populations, which would provide evidence for the evolutionary potential of this plastic response.

## Material & Methods

### Study species

*Antirrhinum majus L.* is an hermaphroditic, self-incompatible (Andalo *et al.*, 2010), short-lived perennial, from the Plantaginaceae family. It has a patchy distribution in the eastern half of the Pyrenees and in the Mediterranean region, from Barcelona (Spain) to Montpellier (France) (Khimoun *et al.*, 2013) at altitudes ranging from 0 and 1900 meters above sea level (Andalo *et al.*, 2010). *A. majus* is a species that colonises open habitats, typically limestone or siliceous substrates, and is adept at colonising screes, road banks and stone walls. *A. majus* is characterised by a wide variation of floral morphology and colour and is easy to grow, with cultivated varieties being widely used in horticulture. It has a diploid genome and has become an important model in population genetics, speciation studies, and floral development research (Schwarz-Sommer *et al.*, 2003).

### Origin of study plants

The plants used in this experiment originate from four natural populations of *A. majus pseudomajus* (Khimoun *et al.*, 2011): Bages (BAG), Banyuls-sur-mer (BAN), Lagrasse (LAG), and Besalù (BES), which are located in Southern France and North Eastern Spain (Supplementary file 1). The general habitat of these four populations is typical of Mediterranean garrigue. In this dry environment, plants grow in a large variety of microhabitats, including open and shaded areas.

The four populations used in this study were chosen on the basis of their high environmental heterogeneity in vegetation cover, resulting in different light conditions within a population. This choice was motivated by the expectation that plasticity is more likely to occur in heterogeneous environments. Additionally, these four populations are significantly genetically differentiated at neutral markers (FST ranging from 0.08 to 0.12, Pujol *et al.*, 2017) and thus can be considered as population replicates within the species level. Seeds were collected on 30 to 50 mature individuals in each population in October 2011.

### Maternal generation in a common environment

In order to reduce the influence of parental environmental effects, we grew a first generation of the seeds in a common environment during spring 2012 in the greenhouse at the CNRS experimental station in Moulis (Station d’Ecologie Théorique et Expérimentale, Moulis, France). We performed within-population crosses during spring and summer 2012 (BAG: 24 mothers, BAN: 25 mothers, BES: 23 mothers, LAG: 15 mothers). This design produced mostly full sib arrays, and some half sib arrays that shared either a father or a mother. Mature seeds were collected during autumn 2012 and were stored in paper bags at room temperature. Subsequently, we used the seeds produced in this common environment to perform a further common garden experiment where we manipulated shade levels.

### Common garden experiment

In June 2014, we sowed seeds from 14 to 16 full sib families (Supplementary file 1) in peat based universal compost in 8*8*7 cm pots. Pots were spatially randomized in the EDB laboratory experimental garden at the Ecole Nationale Supérieure de Formation de l’Enseignement Agricole (ENSFEA, Toulouse-Auzeville). The bases of the pots rested on an irrigation mat (400g/m^2^) to regulate water. A layer of soil of a couple of centimetres covered the bases of the pots and the irrigation mat. To limit neighbour interference, pots were separated by at least 20 cm from each other. All 336 plants that germinated survived, of which 88 individuals flowered. From June until September, approximately half the plants (*n* = 169) were exposed to natural light. The other half of the plants (*n* = 167) were exposed to a shade treatment. Shade was created for each plant individually, using vertical tunnels (height = 40cm) that filtered the light that could reach the plant. These tunnels were made of a metallic mesh shaped into a tube, which was covered in green shading cloth, with an open top. All plants were watered similarly, with no nutrient addition, only when the lack of natural rain made it necessary to provide water to ensure the survival of plants in small pots. Average monthly temperatures ranged from 20.6 to 21.5°C, and cumulative monthly rainfall ranged from 28.3 to 73.4 mm.

The shade treatment modified the light spectrum that the plants received (Supplementary file 2). In particular, it reduced the levels of Photosynthetically Active Radiation (PAR). PAR measurements were taken once on a sunny day with a spectrometer (model AVASPEC-ULS 2048-USB2, Avantes, Apeldoorn, the Netherlands), from 8 am to 12 am, with one measurement per minute. The average PAR in the light treatment was 390.6 μmol.m^-2^.s^-1^, and its average in the shade treatment was 107.2 μmol.m^-2^.s^-1^. The average Red to Far Red ratio (R/FR) was reduced from 2.4 in the light to 1.7 in the shade, as expected from the effect of a green shading cloth.

### Trait measurements

In order to quantify the shade avoidance response in the different populations, we took measures of plant height, internode length, and specific leaf area (SLA). We measured the vegetative height of every plant 40 days after germination, measured as distance (in centimetres) between the ground level and the last node at the top of the plant, or the last node below the first inflorescence if an individual flowered. We counted the number of nodes on this part of the stem, and divided the height by the number of nodes to obtain the mean internode length of 40 day old plants (*Internode 40 days*, available for 327 individuals).

A similar measure was recorded at the date of first flowering (*Internode first flower*). However, flowering inhibition in the shade treatment resulted in very unbalanced data for this trait (N = 67 in the light and N = 21 in the shade, half of the latter belonging to the LAG population). We therefore chose not to perform analysis on this measurement other than testing for its correlation to the other measurements of internode length (see below). In order to obtain a measure of internode length for every individual, we measured the average internode length of the first six nodes of the stem (*Internode six nodes*), on both flowering and non flowering plants at the end of the experiment (N = 336).

For every plant, we estimated the specific leaf area (*SLA*), which is the ratio of the leaf area to its dry mass. For this, we sampled five mature non-senescent leaves on each plant. These leaves were stored in water and in the dark during 6h to 8h before being scanned. They were then kept in silica gel until drying in the oven at 65°C for three days before weighing. SLA was derived by taking the ratio of the area of leaves to the mass of the dried leaves (cm^2^.g^-1^).

Finally, at the end of the experiment, we estimated the total height of each plant, from the ground level to the top (*Height*, which includes the height of the inflorescence in individuals that flowered), and recorded whether each plant had flowered (*Flowering*).

### Phenotypic plasticity

There are many ways to quantify phenotypic plasticity. From a statistical perspective, plasticity can be estimated by the slope of the reaction norm (i.e. the change in phenotypic trait value) for the same genotype in two or more environments. When the same genotype cannot be assessed in several environments (such as in our case), a common method is to estimate the average phenotypic response in the two environments for members of the same family, as they share a similar genetic background. Several metrics have been used in the literature to characterise reaction norms (Valladares *et al.*, 2006). We chose to present a modification of a simple and robust metric, the plasticity index (Valladares *et al.*, 2006), which is estimated as 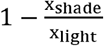 when the value is larger in the light than in the shade, and 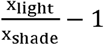 otherwise, where Xshade is the trait value in the light and xshade is the trait value in the shade. Contrary to the original metric (which would be 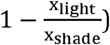, this index ranged between −1 and 1, with values less than zero representing a relative increase of the trait in the shade compared to the light treatment. This index was estimated separately for each family, and at the population scale, using the mean family trait values to reduce the unequal contribution of families to the overall index.

### Statistical analysis

We first estimated the correlation between our three measurements of internode length (*Internode 40 days, Internode first flower* and *Internode six nodes*), by using a Pearson correlation test, to evaluate whether these traits were providing us with similar information. As these measures were well correlated, we only present analysis involving *Internode six nodes*, for which we had most data. We then estimated the correlations between *Height* and *Internode six nodes, Internode six nodes* and *SLA*, and between *Height* and *SLA*.

We used a Generalized Linear Mixed Model approach to estimate the effect of the population of origin (*Population*) and the shade treatment (*Shade*) on *Height, SLA*, and *Internode six nodes*. We ran two models that addressed slightly different questions. The first model estimated the variation in the plastic response as the variation of family slopes between the two treatments (*i.e.* magnitude and direction of the change in trait values), and quantified approximately the genetic variance underlying the response. This approach in effect treats plasticity as a trait in itself (e.g. Scheiner and Lyman, 1989). The second model considers the phenotypes expressed in the different environments as different traits, and estimates the genetic covariance or correlation between the two states (following the approach by Via and Lande, 1985).

### Model 1: Families-level reaction norms

The first model built on the fact that we had measurements for several maternal family members in each treatment, which allowed us to estimate the between-family variance of the slope of the reaction norm between the light and the shade environments. This model included *Shade* and *Population* as fixed factors, as well as their interactions. We also included the interaction *Mother:Shade* as a random effect, which allowed us to test whether individuals from different families had different reaction norms. We allowed different residual variances for light and shade treatments. This model provided estimates for the trait variance in the light treatment, the variance of the slope of the reaction norm between light and shade, as well as the covariance between the intercept in the light treatment and the slope. From these three estimates, we manually derived estimates and confidence intervals for the between-family variance for the trait in the two levels of the *Shade* treatment.

### Model 2: Animal model

The second model is a quantitative genetic Animal Model, which fully accounts for the different degrees of relatedness between individuals in our dataset (*i.e.* full sibs and half sibs). The added value of this approach compared to a classical linear mixed model is that it uses the relationship between individuals separated by different degrees of relatedness to estimate breeding values and thus the additive genetic component of the trait (the *Animal* random factor in our model). Our model included the same fixed effects as the first model: *Shade, Population* and their interaction, and *Animal:Shade* as random effect. We used an unstructured variance-covariance matrix for the interaction between *Animal* and *Shade*, in order to estimate the additive genetic variance in both treatments, as well as the genetic covariance between the two treatments. From this we derived genetic correlations between the trait value in the light environment and the trait value in the shade environment for each trait (Falconer and Mackay, 1996). This genetic correlation shows how the expression of the phenotype is affected across the two environments. Finally, a heterogeneous residual variance was included for each shade treatment, allowing different residual variances in the light and shade, but no covariance between the residuals in the two treatments.

The models were fitted in a Bayesian framework using MCMCglmm (Hadfield, 2016) in the statistical program R (R Core Team, 2016). Priors followed an inverse-wishart distribution (V = diag(2) and nu = 3 for the *Animal:Shade* and the *Mother:Shade*, and V = diag(2) and nu = 0.002 for the residuals). We used a burn-in time of 900000, and 4800000 iterations, with a thinning of 3000, which allowed us to sample the posterior distribution 1300 times (except for *Internode six nodes,* were we used 1500000 burning iterations, 8000000 iterations and 5000 thinning to reduce autocorrelation). We used posterior distributions and model autocorrelation to assess the quality of model runs. Finally, we ran identical models with identical priors five times and used the Gelman-Rubin convergence criterion to assess run quality. This step yielded identical results, demonstrating high reliability between runs. 95% Credible Intervals (CIs) were derived from the runs for all parameters. Effects were considered significant if their 95% credible intervals did not overlap 0.

## Results

### Trait correlations

The three measures of internode length were strongly positively correlated (*Internode six nodes* and *Internode 40 days*: ρ = 0.60, p < 10^−15^, *Internode six nodes* and *Internode first flower*: ρ = 0.73, p < 10^−15^, *Internode 40 days* and *Internode first flower*: ρ = 0.57, p < 10^−7^). We thus chose to focus on only one of these measurements, the *Internode six nodes* for the following analyses. This measure had a strong positive correlation with *Height* (ρ = 0.7, p < 10^−16^). *SLA* was correlated to *Internode six nodes* (ρ = 0.52, p < 10^−16^) and significantly but weakly correlated to *Height* (ρ = 0.23, p < 10^−4^).

### Trait means

Overall, mean *Height* was 29.4 cm (sd = 16.2 cm) and was 23% to 42% less in the light than in the shade depending on the population (Figure 1). Overall, mean internode length (*Internode six nodes*) was 2.6 cm (sd = 1.1 cm) and was, depending on the population, 42% to 47% shorter in the light than in the shade (Figure 2). Overall, average *SLA* was 285.8 cm^2^.g^-1^ (sd = 158.1 cm^2^.g^-1^) and was 63% smaller in the light than in the shade (Figure 3). The probability of *Flowering* was 53% to 85% less frequent in the shade than in the light. We did not conduct an in-depth statistical analysis of this trait because few plants flowered, which would have led to unbalanced analyses.

**Figure 1.**
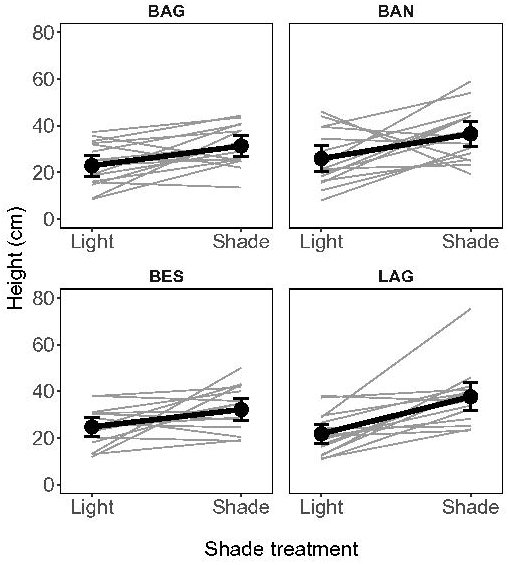
Longer stems in the shade than in the light. Grey: family means. Black dots and lines: population means based on family means. Error bars represent 95% credible intervals.

**Figure 2.**
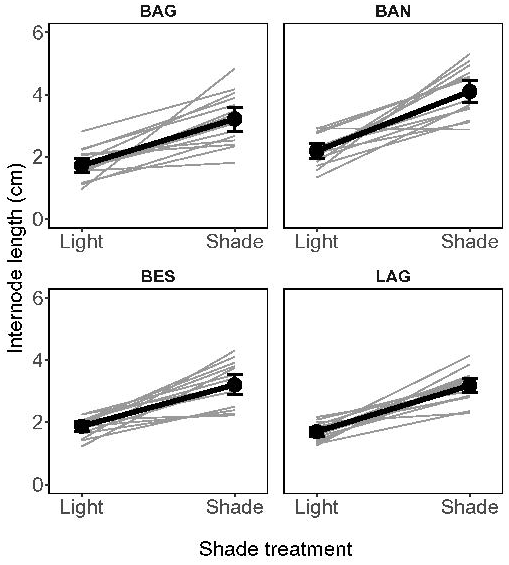
Longer internode length in the shade than in the light. Grey: family means. Black dots and lines: population means based on family means. Error bars represent 95% credible intervals.

**Figure 3.**
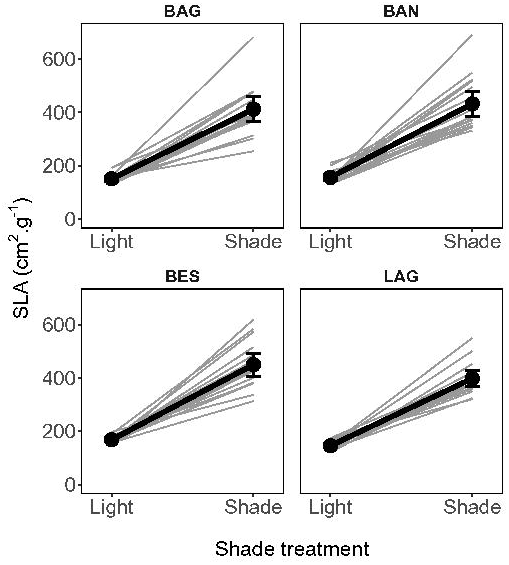
Higher specific leaf area in the shade than in the light. Grey: family means. Black dots and lines: population means based on family means. Error bars represent 95% credible intervals.

### Model 1

The effect of *Shade* on *Height* was significant (Table 2, *Light*: 20.9 cm (CI: 17.3 – 23.6); *Shade*: 31.2 cm (CI: 26.1 – 35.8)). No differences in mean *Height* were detected among the four populations (Table 2, Supplementary file 3). There was no interaction between *Population* and the *Shade* treatment, as shown by null parameters for the interaction term (Supplementary file 4). There was significant variance between families for the intercept, *i.e.* mean trait values differed among families in the light treatment (V_Light_ = 0.14 (CI: 0.09 – 0.27), V_Shade_ = 0.25 (CI: 0.15 – 0.37), Table 3). Families also significantly differed in their response to the shade treatment: there was significant variance between families for the slope of the reaction norm (V_slope_ = 0.17 (CI: 0.1 – 0.31), Table 3). The covariance between the intercept (in the light) and the slope of the reaction norm was not different from zero (cov = 0.02 (CI: −0.11 – 0.14), Table 3).

**Table 1.** Phenotypic trait values in the light and in the shade, by population. Mean (sd) of populations for the traits in all populations, in both the light and shade environments.

| Trait | Population | Light | Shade | Plasticity index |
| --- | --- | --- | --- | --- |
| **Flowering** | BAG | 0.4 (0.3) | 0.1 (0.1) | 0.85 |
|  | BAN | 0.3 (0.3) | 0 (0.1) | 0.92 |
|  | BES | 0.4 (0.3) | 0.1 (0.2) | 0.68 |
|  | LAG | 0.4 (0.3) | 0.2 (0.3) | 0.53 |
| **Height**<br>(cm) | BAG | 22.8 (9.3) | 31.3 (8.8) | -0.27 |
|  | BAN | 25.9 (11.6) | 36.5 (11.3) | -0.29 |
|  | BES | 24.7 (8.4) | 32.1 (9.8) | -0.23 |
|  | LAG | 21.7 (8.6) | 37.6 (12.2) | -0.42 |
| **Internode**<br>**six nodes**<br>(cm) | BAG | 1.7 (0.5) | 3.2 (0.8) | -0.46 |
|  | BAN | 2.2 (0.5) | 4.1 (0.7) | -0.47 |
|  | BES | 1.9 (0.3) | 3.2 (0.7) | -0.42 |
|  | LAG | 1.7 (0.3) | 3.2 (0.5) | -0.47 |
| **SLA**<br>(cm<sup>2</sup>.g<sup>-1</sup>) | BAG | 151 (18.2) | 411.9 (98.4) | -0.63 |
|  | BAN | 156.1 (24) | 431.7 (98.7) | -0.64 |
|  | BES | 167.9 (14.5) | 449.5 (88.6) | -0.63 |
|  | LAG | 145.6 (15.9) | 397.3 (60.6) | -0.63 |

**Table 2.** Significance tests for the *Population* and the *Shade* treatment. Chi-square tests are given for the two models and the three traits of interest (*Height, Internode six nodes*, and *SLA*).

| **Trait** | **model** | **D.I.C.** | **Treatment** | **Chi<sup>2</sup>** | **D.f.** | **P.** |
| --- | --- | --- | --- | --- | --- | --- |
| ***Log Height*** | Model 1 | 604.2 | *Population* | 1.5 | 6 | 0.96 |
|  |  |  | *Shade* | 22.7 | 4 | **0.00** |
|  | Model 2 | 282.8 | *Population* | 1.3 | 6 | 0.97 |
|  |  |  | *Shade* | 15.3 | 4 | **0.00** |
| ***Internode 6 nodes*** | Model 1 | 754.6 | *Population* | 15.8 | 6 | **0.01** |
|  |  |  | *Shade* | 165.7 | 4 | **<10<sup>-3</sup>** |
|  | Model 2 | 475.8 | *Population* | 15.7 | 6 | **0.02** |
|  |  |  | *Shade* | 114.0 | 4 | **<10<sup>-3</sup>** |
| ***Log SLA*** | Model 1 | -55.2 | *Population* | 1.9 | 6 | 0.93 |
|  |  |  | *Shade* | 314.9 | 4 | **<10<sup>-3</sup>** |
|  | Model 2 | -898.2 | *Population* | 4.9 | 6 | 0.55 |
|  |  |  | *Shade* | 304.0 | 4 | **<10<sup>-3</sup>** |

D.f: degrees of freedom; P: P-value.

**Table 3.**
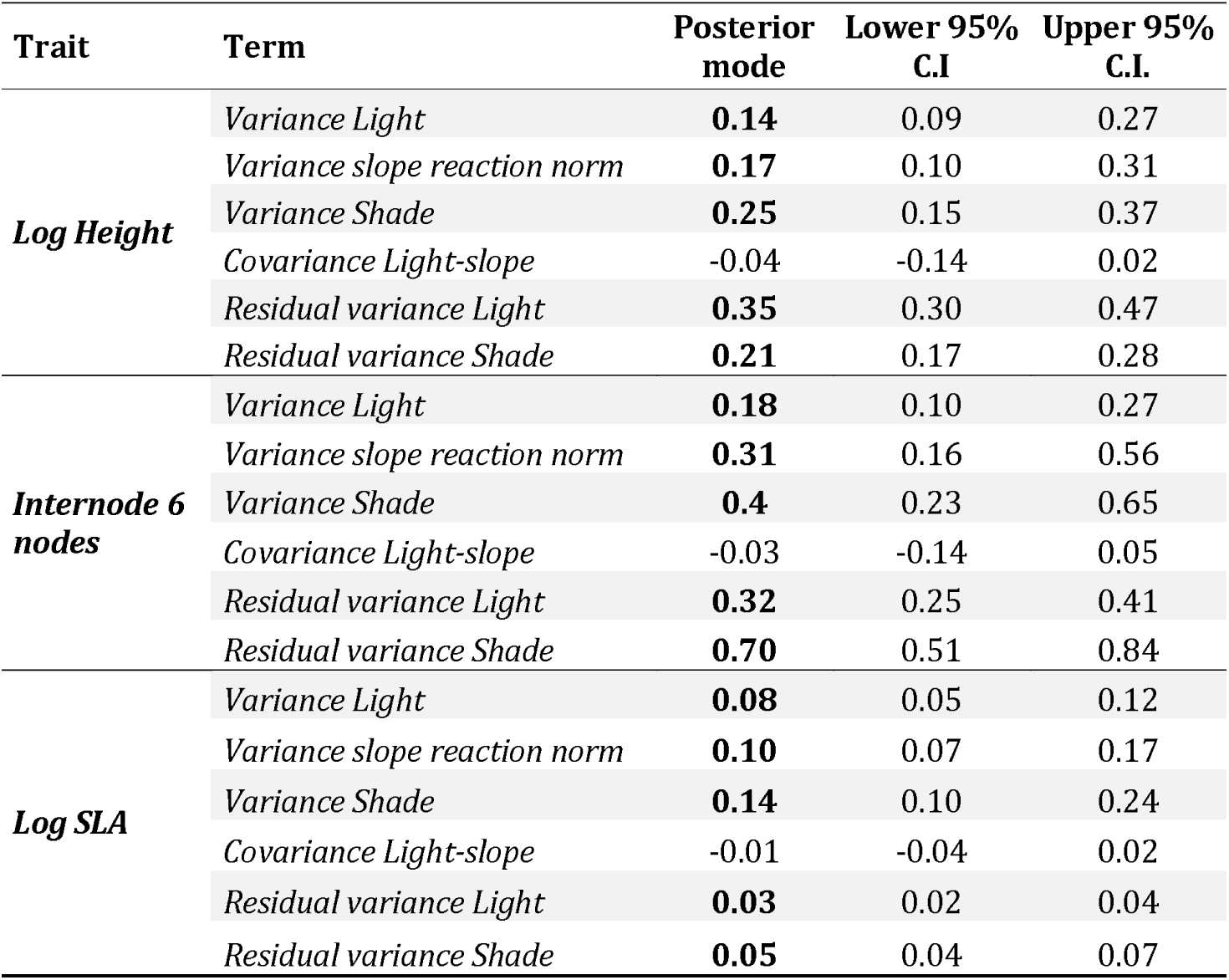
Family variation in the intercept and the slope of the reaction norm (Model 1). Parameter estimates for the random effects estimated by Model 1 are given by posterior mode estimates and their 95% credible intervals (C.I.) for the three traits of interest (*Height, Internode six nodes,* and *SLA*). The last two traits were log transformed to meet the assumptions of the linear model. Values that were considered significant after examination of their credible intervals are presented in bold.

| Trait | Term | Posterior mode | Lower 95% C.I | Upper 95% C.I. |
| --- | --- | --- | --- | --- |
| **Log Height** | *Variance Light* | **0.14** | 0.09 | 0.27 |
|  | *Variance slope reaction norm* | **0.17** | 0.10 | 0.31 |
|  | *Variance Shade* | **0.25** | 0.15 | 0.37 |
|  | *Covariance Light-slope* | -0.04 | -0.14 | 0.02 |
|  | *Residual variance Light* | **0.35** | 0.30 | 0.47 |
|  | *Residual variance Shade* | **0.21** | 0.17 | 0.28 |
| **Internode 6 nodes** | *Variance Light* | **0.18** | 0.10 | 0.27 |
|  | *Variance slope reaction norm* | **0.31** | 0.16 | 0.56 |
|  | *Variance Shade* | **0.4** | 0.23 | 0.65 |
|  | *Covariance Light-slope* | -0.03 | -0.14 | 0.05 |
|  | *Residual variance Light* | **0.32** | 0.25 | 0.41 |
|  | *Residual variance Shade* | **0.70** | 0.51 | 0.84 |
| **Log SLA** | *Variance Light* | **0.08** | 0.05 | 0.12 |
|  | *Variance slope reaction norm* | **0.10** | 0.07 | 0.17 |
|  | *Variance Shade* | **0.14** | 0.10 | 0.24 |
|  | *Covariance Light-slope* | -0.01 | -0.04 | 0.02 |
|  | *Residual variance Light* | **0.03** | 0.02 | 0.04 |
|  | *Residual variance Shade* | **0.05** | 0.04 | 0.07 |

The effect of *Shade* on *Internode six nodes* was significant (Table 2, *Light*: 1.9 cm (CI: 1.8 – 2); *Shade*: 3.4 cm (CI: 3.2 – 3.6)). There were significant differences in average internode length among populations (Table 2, Supplementary file 3), but no interaction between *Population* and *Shade* (Supplementary file 4). There was significant variance between families for the intercept (V_Light_ = 0.18 (CI: 0.10 – 0.27), V_Shade_ = 0.4 (CI: 0.23 – 0.65), Table 3). There was significant variance between families for the slope of the reaction norm (V_slope_ = 0.31 (CI: 0.16 – 0.56), Table 3). The covariance between the intercept and the slope of the reaction norm was not different from zero (cov = −0.04 (CI: −0.14 – 0.02), Table 3).

The effect of *Shade* on *SLA* was significant (Table 2, *Light*: 153.4 m^2^.g^-1^ (CI: 139.5 – 165.6); *Shade*: 412.5 cm^2^.g^-1^ (CI: 357.6 – 459.4)). No differences in average *SLA* were detected among the four populations (Table 2, Supplementary file 3). There was no interaction between *Population* and the *Shade* treatment (Supplementary file 4). There was significant variance between families for the intercept (V_Light_ = 0.08 (CI: 0.05 – 0.12), V_Shade_ = 0.14 (CI: 0.1 – 0.24), Table 3) and for the slope of the reaction norm (V_slope_ = 0.1 (CI: 0.07 – 0.17), Table 3). The covariance between the intercept and the slope was not different from zero (cov = −0.01 (CI: −0.04 – 0.02), Table 3).

In summary, all studied populations of *A. majus* exhibited a strong response to shade for all the three traits. There was no evidence of an interaction between *Population* and *Shade* in either of the traits (Supplementary file 3). The significant variance for families intercept and slope for the three traits meant that the plastic responses of different families differed in magnitude and direction, despite the fact that all individuals shared the same maternal environment. The null covariance between the intercept and slope of the reaction norm indicated that at the family level, the slope of the plastic response was independent of the mean trait value in the light treatment. In other words, families that were on average taller in the light did not always have a stronger plastic response than families that were on average shorter in the light.

### Model 2

The effect of *Shade* on *Height* was significant (Table 2, *Light*: 20.6 cm (CI: 17.3 – 23.8); *Shade*: 30.2 cm (CI: 26.6 – 36)). No differences in mean *Height* were detected among the four populations (Table 2, Supplementary file 3). There was additive genetic variance in both light (V_Light_ = 0.24 (CI: 0.12 – 0.54), Table 4) and shade (V_Shade_ = 0.28 (CI: 0.15 – 0.43), Table 4). The genetic covariance of the trait in the *Light* and *Shade* treatment was null, and with large credible intervals for (Table 4).

**Table 4.**
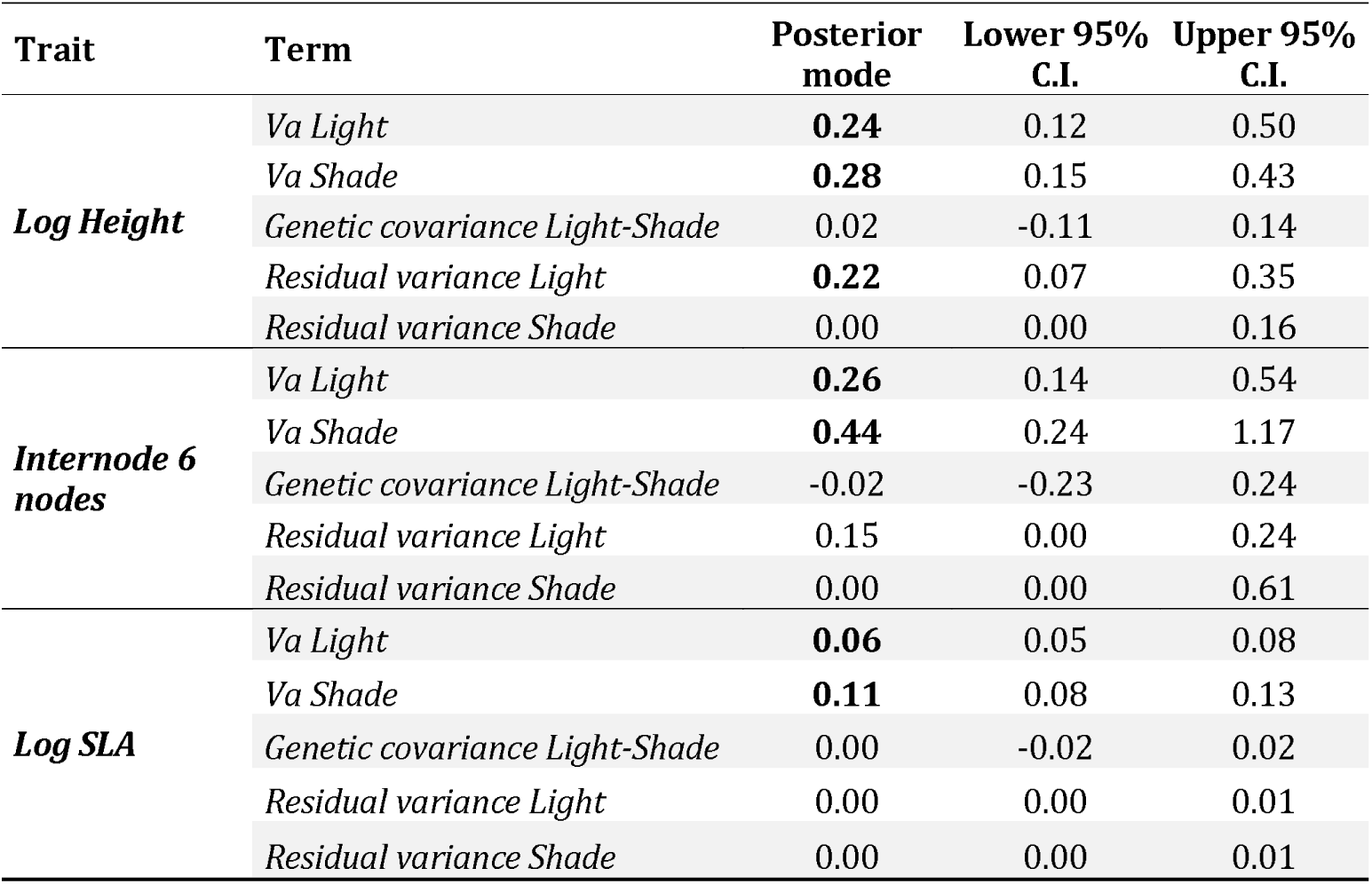
Model 2 Genetic additive variance and covariances between light and shade (Model 2). Parameter estimates for the random effects estimated by Model 2 are given by posterior mode estimates and their 95% credible intervals (C.I.) for the three traits of interest (*Height, Internode six nodes, and SLA*). The last two traits were log transformed to meet the assumptions of the linear model. Values that were considered significant after examination of their CI are presented in bold.

| Trait | Term | Posterior mode | Lower 95% C.I. | Upper 95% C.I. |
| --- | --- | --- | --- | --- |
| **Log Height** | *Va Light* | **0.24** | 0.12 | 0.50 |
|  | *Va Shade* | **0.28** | 0.15 | 0.43 |
|  | *Genetic covariance Light-Shade* | 0.02 | -0.11 | 0.14 |
|  | *Residual variance Light* | **0.22** | 0.07 | 0.35 |
|  | *Residual variance Shade* | 0.00 | 0.00 | 0.16 |
| **Internode 6 nodes** | *Va Light* | **0.26** | 0.14 | 0.54 |
|  | *Va Shade* | **0.44** | 0.24 | 1.17 |
|  | *Genetic covariance Light-Shade* | -0.02 | -0.23 | 0.24 |
|  | *Residual variance Light* | 0.15 | 0.00 | 0.24 |
|  | *Residual variance Shade* | 0.00 | 0.00 | 0.61 |
| **Log SLA** | *Va Light* | **0.06** | 0.05 | 0.08 |
|  | *Va Shade* | **0.11** | 0.08 | 0.13 |
|  | *Genetic covariance Light-Shade* | 0.00 | -0.02 | 0.02 |
|  | *Residual variance Light* | 0.00 | 0.00 | 0.01 |
|  | *Residual variance Shade* | 0.00 | 0.00 | 0.01 |

This effect of *Shade* on *Internode six nodes* was significant (Table 2, *Light*: 1.9 cm (CI: 1.7 – 2.1); *Shade*: 3.4 cm (CI: 3.2 – 3.7)). There were significant differences in average internode length among populations (Table 2, Supplementary file S3). There was additive genetic variance in both light (V_Light_ = 0.26 (CI: 0.14 – 0.54), Table 4) and shade (V_Shade_ = 0.44 (CI: 0.24 – 1.17), Table 4), but no genetic covariance of the trait between the *Light* and *Shade* treatment (Table 4).

The effect of *Shade* on *SLA* was significant (Table 2, *Light*: 151.1 cm^2^.g^-1^ (CI: 141 – 161.3); *Shade*: 340.3 cm^2^.g^-1^ (CI: 373.6 – 445.6)). No differences in average *SLA* were detected among the four populations (Table 2, Supplementary file 3). There was additive genetic variance in both light (V_Light_ = 0.06 (CI: 0.05 – 0.08), Table 4) and shade (V_Shade_ = 0.11 (CI: 0.08 – 0.13), Table 4), but no genetic covariance of the trait between the *Light* and *Shade* treatment (Table 4).

In summary, all studied populations reacted strongly to Shade, as in Model 1. Model 2 estimated null (and small) covariances between the *Light* and *Shade* treatment, which means that individuals with high trait values in the light did not consistently have high (or low) trait values in the shade environment. Consequently, the very weak genetic correlations that derive from these covariances had very large credible intervals (*Log Height*: 0.16 (CI:−0.32 – 0.49); *Internode six nodes*: 0.10 (CI:-0.44 – 0.45); *Log Log SLA*: −0.02 (CI:-0.25 – 0.21)).

## Discussion

Our results showed that *A. majus* plants have a large phenotypic plastic response to shade. When exposed to shade, *A. majus* plants increased in height, internode length, and SLA, typical of the shade avoidance syndrome (Smith and Whitelam, 1997). In a situation where light may become, or is already, scarce, stem elongation through increased distance between nodes and/or stem height is expected to allow plants that are not shade tolerant to outgrow competitors. An increase in SLA in the shade, while technically not a shade avoidance strategy, is a functional response that has often been observed (Lewandowska and Jarvis, 1977; Dong, 1995; Evans and Poorter, 2001; van Kleunen *et al.*, 2011; Liu *et al.*, 2016), and is expected to improve the physiological performance of plants in a light-limited environment.

Similar plastic responses have been observed in other studies involving *A. majus* or other *Antirrhinum* cultivars. In *A. majus* cultivars, Munir et al. (2004) found increased total height and reduced branching at lower light intensity and Khattak & Pearson (2005) found higher internode length in reaction to light filtering. In addition, Cremer et al. (1998) observed shorter flowering time under low R/FR ratio in an inbred *A. majus* line (Sippe50). Our results demonstrate the existence of this syndrome in *A. majus* plants from wild populations. Furthermore, we did not observe different levels of plasticity in the four populations. This does not necessarily mean that plants from these populations will express the same responses when growing in wild conditions. Complex environments may constrain the expression of plasticity, particularly if some phenotypes are more costly than others in a particular set of conditions (Donohue *et al.*, 2000; Huber *et al.*, 2004). The plastic response to shade is an evolutionary labile trait that can be lost quickly during evolutionary transitions, even at the within-species scale, such as between wild populations and cultivars (Pujol *et al.*, 2005). However, the combined results from past studies of *A. majus* cultivars and this study support the hypothesis that plasticity is widespread among *A. majus* populations, whether it evolved post-speciation or was conserved from ancestral species.

For such a set of plastic responses to evolve, or to be conserved, individuals need to have access to reliable cues that describe the current and/or future light environment. Plants usually rely on a reduction of the ratio of red to far-red radiations (R/FR ratio) as a cue indicating the presence of competitors for light: plant responses under reduced R/FR ratios are similar to those observed in dense patches where there is competition for light (Smith, 1982; Ballaré *et al.*, 1987, 1990). The reduction of total light intensity, and in particular blue light, also induces a shade avoidance response in some species (Ballaré, 1999; Pierik *et al.*, 2004; Franklin, 2008, 2016; Ballaré and Pierik, 2017). Previous studies on *A. majus* inbred lines and cultivars showed that the reduction of both the R/FR ratio and the amount of blue radiation are likely to be involved in *A. majus* shade avoidance response (Cremer *et al.*, 1998; Khattak and Pearson, 2005). However, no study had evaluated this plastic response in *A. majus* plants originating from natural populations. In our experiment, we did not attempt to test these two cues in isolation. Instead we evaluated the effect of a controlled light environment simulating competition but excluding other effects of high density (e.g., plant hormones modification, nutrient or water competition), and found that *A. majus* responded strongly to this controlled light environment, suggesting that either reduction of light intensity, R/RF ratio or both act as cues in this species.

We performed a study of the population variation of shade-avoidance responses, both between populations and between families, for several morphological and functional traits. Our results showed that families within populations varied in their phenotypic plasticity to shade, thereby supporting the presence of a genetic basis for this plasticity. Family effects are often used to evaluate the genetic component of traits because members of the same family are genetically related (Lynch and Walsh, 1998). This effect must be interpreted in the broad sense, however, because it encompasses additive genetic, dominance, maternal, and epistatic effects. While growing the parental generation in a common environment reduces the possible environmental component of maternal effects, it does not entirely eliminate it, and we cannot distinguish between all components of variance with our design.

Genotype by environment interactions are often found in studies of phenotypic plasticity (Pigliucci, 2005). However, only a few cases of GxE interactions were documented for the plastic response of plants to shade or competition. For example, Donohue and Schmitt (1999) found a G×E interaction in the plastic response to plant density in natural settings, using inbred lines of *Impatiens capensis*. Botto & Smith (2002) found a G×E interaction for the plastic response of *Arabidopsis thaliana* accessions to R/FR ratio modifications. However, for the shade avoidance response, most known G×E interactions correspond to comparisons between a plant wild type and mutant lines (Skalova and Krahulec, 1992; Andersson and Shaw, 1994; Schmitt *et al.*, 1995, 2003), or between populations (Pigliucci *et al.*, 1995; Van Hinsberg, 1997). Results from past studies did not necessarily support family variation in the plastic response to shade, i.e. variation in the slope of the reaction norm in plants from natural populations (e.g. Andersson and Widén, 1994). The genetic variation in plasticity that we found is not surprising, as the molecular pathways are extremely complex (e.g. Franklin and Whitelam, 2005; Vandenbussche *et al.*, 2005; Franklin, 2008; Pierik *et al.*, 2013; Ballaré and Pierik, 2017). It is therefore logical to expect that the great number of genes involved in these pathways would contribute to within-population variation. As this genetic variation in the plastic response is the basis for the evolution of different degrees of plasticity, our results imply that wild populations of *A. majus* have the potential to evolve under selection for higher or lower plasticity.

Additionally, we investigated the trait genetic covariances between the two environments for our three traits, which provides us with information on the constraints acting on the evolution of plasticity (Via and Lande, 1985; van Kleunen and Fischer, 2005; Pujol *et al.*, 2018). We found no genetic covariances between trait values measured in the light and in the shade treatment, in all traits. Furthermore, the family means in the light treatment did not strongly co-vary with the slope of the reaction norm, which suggests no genetic correlations between the trait value in the light treatment and the plastic response. Collectively, these results suggest that, at least in experimental conditions, there were few genetic constraints on the expression of the trait in the light and in the shade. These results imply that the genes involved in the mean trait response in one environment are partially independent from the genes acting on the plastic response. This could be that the gene expressed in the light environment are mostly different than the genes expressed in the shade environment, but more likely this implies the existence of regulator genes which affects the expression of the same genes differently in the light and in the shade, thus mediating the effects of the light spectral cues. In contrast, other studies that estimated the genetic covariance or correlations between traits under light and shade conditions found a much higher correlation between the mean of the trait and the magnitude of the plastic response (Donohue and Schmitt, 1999; Donohue *et al.*, 2000). The relevance of our findings is obviously limited to their experimental background and it would be useful to evaluate whether our conclusions hold in the more stringent natural settings of the four natural populations that we studied. Nevertheless, our current findings suggest that genetic correlations do not constrain the reaction norm to shade in these *A. majus* populations.

### Conclusion & Perspectives

Our experimental findings imply that *A. majus* plants can react to shade and competition for light in natural populations. This plasticity gives them the potential to respond to the temporal and spatial variation of density. Our results support the presence of genetic variation for the traits under both light and shade environments, between-family variation for the slope of the reaction norm, with no evidence for genetic correlations that could constrain the evolution of plasticity. Collectively, our findings strongly support the hypothesis that the shade avoidance response in *A. majus* is not only widespread, but has the potential to evolve in response to selection.

## Acknowledgments

We thank David Field, Caroline Thomson and Isabel Winney for constructive comments on this manuscript. This project has received funding from the European Research Council (ERC) under the European Union’s horizon 2020 research and innovation program (grant agreement No ERC-CoG-2015-681484-ANGI) awarded to BP. This work was supported by funding from the French “Agence Nationale de la Recherche” (ANR-13-JSV7-0002 “CAPA”) to BP. This project was also supported by the ANR funded French Laboratory of Excellence projects “LABEX TULIP” and “LABEX CEBA” (ANR-10-LABX-41, ANR-10-LABX-25-01).

## Supplementary files

Supplementary information is available at Heredity’s website”

## Data archiving

Data and R statistical analysis protocols used in this article will be submitted to Zenodo and will be made available to the reviewers upon request of the editor.

